# Debates about vaccines and climate change on social media networks: a study in contrasts

**DOI:** 10.1101/2020.11.24.396226

**Authors:** Justin Schonfeld, Edward Qian, Jason Sinn, Jeffery Cheng, Madhur Anand, Chris T. Bauch

**Author notes:** **Materials and correspondence** Chris T. Bauch.

## Abstract

Vaccines and climate change have much in common. In both cases, a scientific consensus contrasts with a divided public opinion. They also exemplify coupled human-environment systems involving common pool resources. Here we used machine learning algorithms to analyze the sentiment of 87 million tweets on climate change and vaccines in order to characterize Twitter user sentiment and the structure of user and community networks. We found that the vaccine conversation was characterized by much less interaction between individuals with differing sentiment toward vaccines. Community-level interactions followed this pattern, showing less interaction between communities of opposite sentiment toward vaccines. Additionally, vaccine community networks were more fragmented and exhibited numerous isolated communities of neutral sentiment. Finally, pro-vaccine individuals overwhelmingly believed in anthropogenic climate change, but the converse was not true. We propose mechanisms that might explain these results, pertaining to how the spatial scale of an environment system can structure human populations.

## Main

Despite a strong scientific consensus that vaccines do not cause autism and that anthropogenic climate change is real, public debate on both topics continues to polarize many populations^1,2^. These public debates are well represented in online social media^3-7^, including Twitter^6,7^. On account of the widespread use of Twitter and the ease through which users may share information, Twitter has been widely used to investigate spread of opinions and sentiment through social networks^8,9^. Echo chambers are also common on Twitter and have been identified in both the climate change debate^10^ and the vaccine debate^11^. Moreover, Twitter communities are often highly divided: a study of Twitter communities and conversations after release of the 2013 Intergovernmental Panel on Climate Change Report found evidence that the discussion on climate change is becoming more polarized over time^12^.

Multiple studies have found that Twitter sentiment is correlated with real-world population behaviour, despite the fact that Twitter users are self-selected and do not represent the general population^13^. For instance, vaccination behaviour surveyed by phone correlates well with vaccination sentiment expressed on Twitter^11^. Twitter has been used to predict and monitor infectious disease outbreaks^14-17^. Emotions communicated on social media are connected to clinical visitations^18^. Both the total volume of market-related tweets and the sentiment of those tweets are correlated to stock market movements^19^. These studies suggest that analyzing public Twitter debates on climate change and vaccines could be helpful for understanding real-world decision-making concerning vaccines and climate change, although such efforts are still in their early stages^8^. As well, the impact of real-world developments in natural systems such as global temperature anomalies are reflected in online social media data^20^.

Despite the surge in research using Twitter to study sentiments and opinions, relatively little work compares widely differing topics such as vaccines and climate change. However, comparing different conversations on Twitter can help to better contextualize any given topic; stimulate new hypotheses about social interactions; and test existing hypotheses. Moreover, the climate change and vaccine conversational topics are more closely connected than they might appear. Both vaccine-generated herd immunity and reduced greenhouse gas emissions represent common pool resources that free-riders can exploit by enjoying the benefits of the common resource without paying for its maintenance^21-23^. Both systems also exemplify coupled human-environment systems, where human dynamics and natural dynamics interact with one another^24,25^, and are highly relevant to United National Sustainable Development Goals^26^. A human-environment approach to sustainability problems is called for^27^, and online social media present opportunities to study how individuals and communities of differing sentiment and opinion interact on these vital issues.

Here we compared user sentiment toward vaccines and anthropogenic climate change in a large database of tweets. We analyzed tweet sentiment with machine learning algorithms^28,29^, inferred the sentiment of users, and compared the structure of user and community networks^30^. Our hypothesis was that both conversations would exhibit qualitatively similar distributions of sentiment and organization of user and community networks. In the following subsections, we analyze the data in order of increasing organizational scale, from tweets, to users, to user networks, and finally to community networks.

## Results

In this section we report results for a dataset encompassing approximately 20 million tweets on climate change and 60 million tweets on vaccines (the “GNIP dataset”, Table 1). Confirmatory results using a smaller second dataset collected through the Twitter REST Application Programming Interface appear in the SI Appendix (the “collected dataset”). Additional details on the search terms and datasets appear in Methods. Unless otherwise specified, results in this section pertain to the GNIP dataset.

**Table 1.** Description of datasets. Counts of the number and type of tweets available in each of the data sets, as well as start and end dates, how many were retweets, and how many included links. ‘M’ denotes millions. Details on search terms appear in Methods.

|  | Tweets | Start Date | End Date | Retweets | Links |
| --- | --- | --- | --- | --- | --- |
| GNIP climate | 18.06 M | 2015-04-27 | 2016-10-26 | 10,16 M (56%) | 12.60 M (70%) |
| GNIP vaccine | 62.11 M | 2006-06-01 | 2016-10-26 | 23.27 M (37%) | 39.00 M (48%) |
| collected climate | 2.71 M | 2016-10-26 | 2018-09-20 | 1.82 M (67%) | 1.60 M (59%) |
| collected vaccine | 4.40 M | 2016-10-26 | 2018-09-20 | 2.24 M (51%) | 2.50 M (57%) |

### Prevalence of sentiment categories in vaccine and climate tweets

We used machine learning algorithms to classify the sentiment of tweets into “pro”, “anti” and “neutral/other” (see Methods for details). A pro-vaccine tweet expressed sentiment that vaccines are safe and effective, while an anti-vaccine tweet expressed sentiment that vaccines are harmful and/or ineffective, or that infectious diseases are not dangerous and do not necessitate vaccination. A neutral vaccine tweet was neither pro-nor anti-vaccine. Similarly, a pro-climate tweet expressed sentiment that human-caused climate change is real and caused by humans; an anti-climate tweet expressed sentiment that climate change is not real, or that humans do not cause climate change, or that climate change is not a problem. And a neutral/other climate tweet was neither pro- nor anti-climate. Neutral tweets often tweeted news article titles without additional context indicating user sentiment about the article, or the sentiment of the tweet was not clear.

The sentiment of tweets in the vaccine and climate conversations were strikingly different (Table 2). The majority of climate change tweets were pro-climate, with fewer neutral or anti-climate tweets. In contrast, a strong majority of tweets about vaccines were neutral, with relatively few pro-vaccine tweets and even fewer anti-vaccine tweets. This was true across three different datasets and classification methods: (1) human classification of 75,000 randomly sampled tweets from the full datasets, and classification of the full datasets using two different machine learning algorithms, either a (2) linear support vector machine or (3) an ensemble method. The linear support vector machine matched the breakdown of tweets observed in the human classified dataset more closely than the ensemble machine learning algorithm, but the ensemble algorithm had a higher accuracy score. Hence we use the ensemble algorithm in the rest of the Results section unless otherwise stated.

**Table 2.** Sentiment of tweets varies between climate change and vaccine discussions. Sentiment breakdown of tweets as assessed by human readers, an ensemble machine learning algorithm, and a support vector machine algorithm. For the human classification dataset 75,000 tweets were drawn at random from the GNIP climate and GNIP vaccine data sets. Each tweet was assigned a sentiment by three independent humans readers. Only tweets where all three humans agreed were used to generate the above breakdown of sentiment. Details on the machine learning algorithms appear in Methods.

| Dataset | Model | Positive | Negative | Neutral |
| --- | --- | --- | --- | --- |
| Climate Training Set | Human classification | 80.53% | 11.26% | 8.21% |
| GNIP Climate | Machine Learning: Ensemble | 53.18% | 9.23% | 37.59% |
| GNIP Climate | Machine Learning: Support Vector Machine | 72.38% | 13.72% | 13.90% |
| Vaccine Training Set | Human classification | 20.67% | 7.75% | 71.58% |
| GNIP Vaccine | Machine Learning: Ensemble | 15.03% | 4.25% | 80.71% |
| GNIP Vaccine | Machine Learning: Support Vector Machine | 19.90% | 10.82% | 69.28% |

### User sentiment and participation in discussions

Next, we moved up to the level of individual Twitter users. We classified users into pro-, anti-, or neutral for both vaccine and climate conversations, based on the sentiment expressed in their tweets and using the same criterion to classify users as pro-, anti-, or neutral (see Methods). We found a very similar breakdown of sentiment between the three categories for users (SI Appendix, Table S1) as we found for the individual tweets (Table 2). The same pattern was also observed when we considered only tweets by moderate users (tweeting 10 times or more on a topic) or high frequency users (tweeting 100 times or more, SI Appendix Table S1). We observed that the collected dataset contained fewer pro-climate tweets than the GNIP dataset, although pro-climate sentiment was still the most common of the three categories (SI Appendix Table S2). The neutral category was even more common in the collected vaccine dataset than the GNIP vaccine dataset (SI Appendix Table S2).

More English-language tweets originate in the United States than in any other country. Comparing these results (Table 2 and SI Appendix Table S2, S3) to surveys of the United States population shows the same pattern whereby more individuals are pro-climate/pro-vaccine than anti-climate/anti-vaccine, as we define them here. For instance, 63% of United States residents believe global warming is happening whereas 16% believe it is not^31^. Similarly, 88% of Americans believe that the benefits of the measles-mumps-rubella vaccine outweighs its risks, while only 10% believe risks outweigh benefits^32^.

Next, we analyzed the sentiment of users who participated in both discussions. Approximately 1.6 million users tweeted at least once in both debates. But only 158,941, 78,058, and 3,980 users tweeted 5, 10, or 100 times or more in both debates, respectively (SI Appendix Table S3). Overall, between 4% and 38% of all users participated in both conversations, depending on the frequency of the user’s tweets and whether the GNIP dataset or the collected dataset was analyzed. This proportion was highest for individuals tweeting 5 or more times in both conversations (SI Appendix Table S3).

We observed a strong asymmetry in participation and user sentiment expressed between the two topics. 38% of users who participated in the climate change discussion also participated in the vaccination discussion. But only 10% of users who participated in the vaccination discussion also participated in the climate change discussion. Among individuals tweeting 100+ times or more in both conversations, pro-vaccine individuals were overwhelmingly pro-climate (98%), although a significant proportion of pro-climate individuals were either anti-vaccine (11%) or neutral (13%) (Table 3). This supports the hypothesis of a liberal bias against vaccines, although the effect is not very strong^33^. This pattern also occurs when analyzing low and moderate frequency users (SI Appendix Table S5, S6). We also observed that users participating in the climate change discussion were more likely to retweet (56% of the time) versus users participating in the discussion on vaccination (37%). Climate change users were more likely to tweet a link than vaccine users.

**Table 3.** Many pro-climate users are anti-vaccine, but the converse is not true. Number of users tweeting in both conversations 100 times or more on each subject, broken down by sentiment, for the GNIP dataset. Differences are statistically significant (*χ*-squared = 308.34, df = 9, *p* < 2.2*−*16).

|  | Vaccine |  |  |  |
| --- | --- | --- | --- | --- |
| Climate | Positive | Neutral | Negative | Divided |
| Positive | 948 | 2663 | 279 | 25 |
| Neutral | 0 | 3 | 0 | 0 |
| Negative | 2 | 16 | 41 | 0 |
| Divided | 1 | 2 | 0 | 0 |

### User networks

Next we moved up to the level of networks of users interacting through retweets and mentions. We considered two general classes of networks: directed and mutual. In a directed retweet network, if A retweeted B and B retweeted A there will be two edges in the resulting network–one from A to B and one from B to A. Mutual means that if A retweeted B and B did not retweet A there would be no edge between A and B; there is only an edge if both A and B retweeted one another. Undirected means there is an edge between A and B if A retweets B or B retweets A, but it does not matter which. Networks for mentions work similarly. The resulting networks–directed, undirected, and mutual retweet and mention networks for both conversations–are large and sparse. For instance, the GNIP vaccine retweet network of low-frequency users (>5 tweets) consists of 1,283,521 nodes and 6,758,282 edges. The GNIP climate retweet network consists of 459,058 nodes and 3,742,314 edges.

A visualization of the mutual retweet and mention without retweet networks for individuals tweeting 100+ times on the topic show a striking contrast between the two conversations (Figure 1). We observe distinct pro-vaccine, anti-vaccine and neutral regions in both retweet and mention vaccine networks. But the climate networks tend to be more pro-climate overall and anti-climate users are dispersed through the network instead of being clumped together, as in the vaccine network. Networks for users who tweeted 5+ or 10+ times were also visualized but are more difficult to interpret on account of the size of the network (SI Appendix Figures S1, S2).

**Figure 1.**
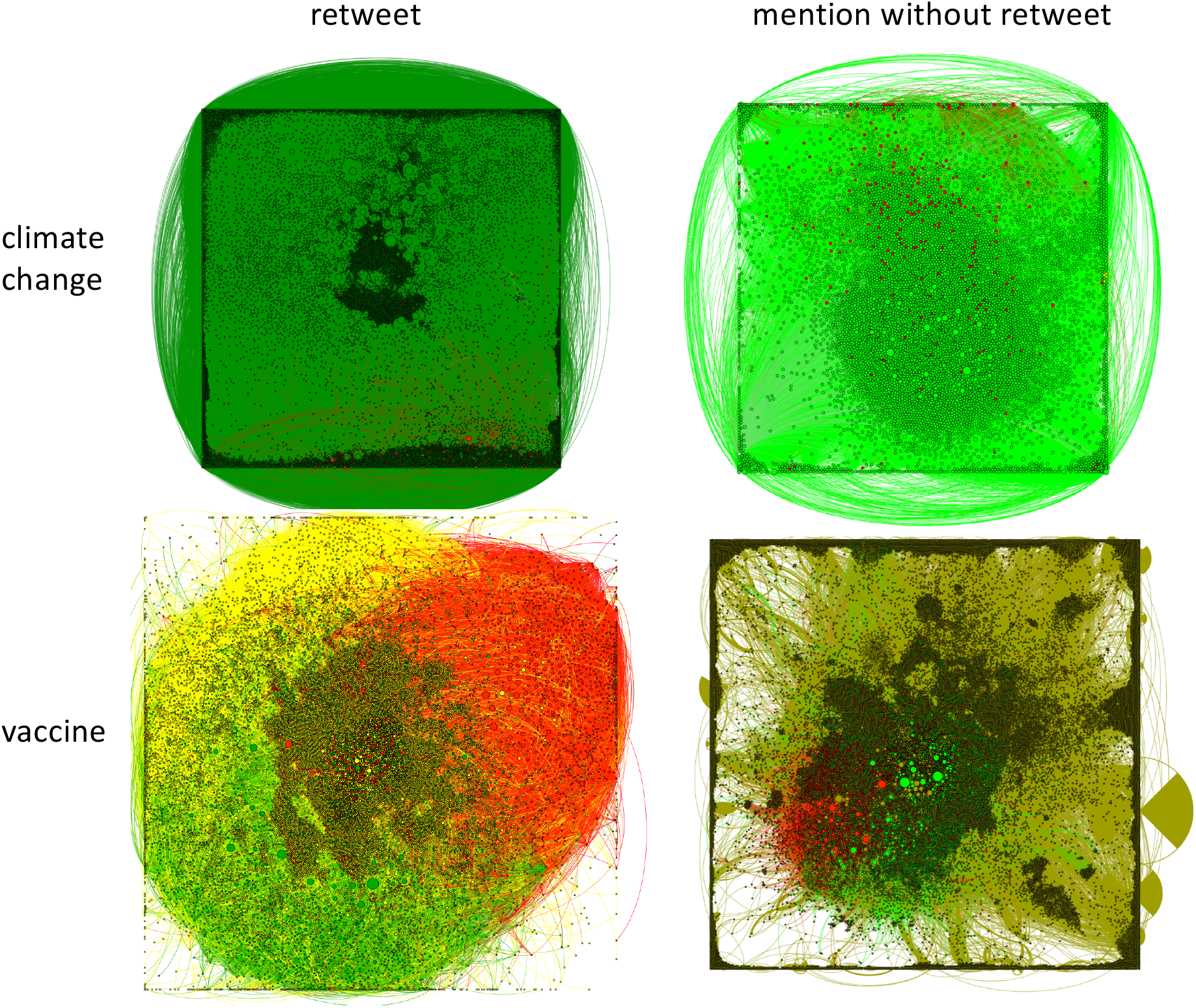
The vaccine conversation is more assortative with respect to sentiment. Network of users tweeting 100 times or more in the climate change (top) and vaccine (bottom) conversations, for retweet (left) and mention (right) networks. Each node represents an individual user. Edges represent interactions between the users: a user has tweeted or mentioned another user. The colour of the node represents the most common sentiment expressed by that user: green represents pro-, red represents anti- and yellow represents neutral.

Network statistics can be computed and confirm the impression that there is less interaction between users of differing sentiment in the vaccine conversation. We computed various network measures, including the assortativity coefficient that measures whether individuals sharing the same sentiment tend to associate with one another more often. (A full list of network statistics for all of our network permutations appears in SI File S1). We found that the vaccine user networks are much more assortative than the climate user networks: users participating in the vaccine discussion tend to retweet and mention individuals who share their sentiment much more than individuals participating in the climate discussion (Table 4).

**Table 4.**
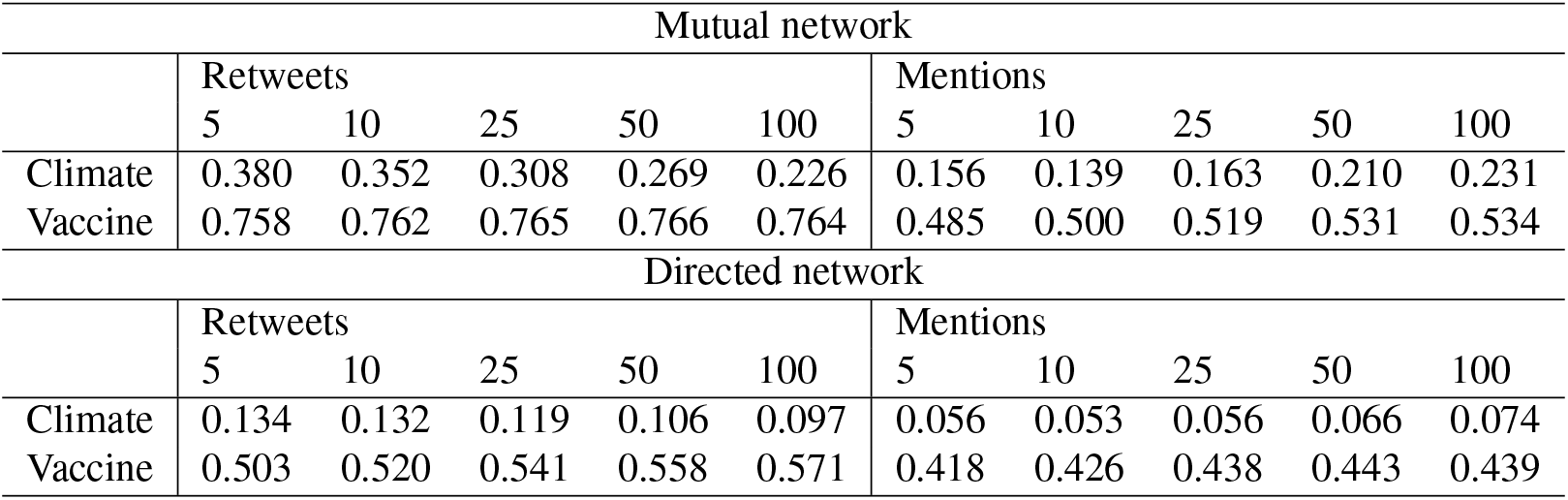
The vaccine conversation exhibits higher levels of assortative mixing. Assortativity coefficients of users according to sentiment using the ensemble machine learning algorithm for tweet sentiment classification and the plurality model for the assignment of user sentiment (see Methods). Mutual network (top) and directed network (bottom). For a P 2-tailed test, *p* « 10*−*4.

| Mutual network |  |  |  |  |  |  |  |  |  |  |
| --- | --- | --- | --- | --- | --- | --- | --- | --- | --- | --- |
|  | Retweets |  |  |  |  | Mentions |  |  |  |  |
|  | 5 | 10 | 25 | 50 | 100 | 5 | 10 | 25 | 50 | 100 |
| Climate | 0.380 | 0.352 | 0.308 | 0.269 | 0.226 | 0.156 | 0.139 | 0.163 | 0.210 | 0.231 |
| Vaccine | 0.758 | 0.762 | 0.765 | 0.766 | 0.764 | 0.485 | 0.500 | 0.519 | 0.531 | 0.534 |
| Directed network |  |  |  |  |  |  |  |  |  |  |
|  | Retweets |  |  |  |  | Mentions |  |  |  |  |
|  | 5 | 10 | 25 | 50 | 100 | 5 | 10 | 25 | 50 | 100 |
| Climate | 0.134 | 0.132 | 0.119 | 0.106 | 0.097 | 0.056 | 0.053 | 0.056 | 0.066 | 0.074 |
| Vaccine | 0.503 | 0.520 | 0.541 | 0.558 | 0.571 | 0.418 | 0.426 | 0.438 | 0.443 | 0.439 |

### Supporting secondary analyses

We carried out various additional analyses to explore the robustness of our finding that the vaccine conversation is more assortative with respect to sentiment. For instance, we found that the vaccine conversation is more assortative in networks of low, moderate, or high frequency users and regardless of whether we consider the directed networks instead of mutual networks (SI File S1). And the pattern persists across a range of permutations for network type and method of classification (SI File S1).

To control for errors in the assignment of tweets to sentiment categories, we considered an 80% threshold approach where an individual would be assigned a sentiment only if at least 80% of their tweets were labeled with that sentiment (In contrast, our baseline analysis used a plurality approach, where a user’s sentiment was considered to be that of whichever type of tweet they tweeted the most: positive, negative, or neutral). The 80% approach has the effect of removing individuals with mixed opinions and/or incorrectly categorized tweets. As expected, this approach generated a higher assortativity across both conversations on account of removing individuals who do not express a clear sentiment in 80% or more of their tweets, many of which might be bots. For both retweets and mentions on the directed network, and for mentions on the mutual network, the assortativity coefficient was higher for the vaccine discussion than the climate discussion. However, for retweets on the mutual network, both discussions exhibited an assortativity coefficient close to 1 (SI Appendix Tables S6, S7).

Our search term list for vaccines was much larger than for climate change (see Methods). This could potential cause different types of information to be sampled. Hence, we explored the effect of subsampling the vaccine data using only three keywords (vaccine, flu, or influenza) and pulling only 28.35% of the data. The assortativity of the subsampled vaccine data set continued to be much higher than that of the climate data set for both mutual and directed networks (SI Appendix Table S8 and S9). The assortativity coefficient was higher in the vaccine conversation when using the collected dataset for both directed networks and for the mutual retweet network, although not for the mutual mention network (SI Appendix Table, S10,S11). Finally, assortativity continued to be higher in vaccine conversation when we used a support vector machine for both the plurality, 80 % threshold, and subsampled cases for the GNIP data (SI Appendix Tables S12-S14).

In total, there were 260 permutations in our analyses, with respect to choice of: vaccine or climate change conversation; retweet versus mention without retweet; subsampling of tweets; using 80% thresholds instead of plurality to assign user sentiment; ensemble versus SVM algorithm; and directed versus mutual network. For each of these permutations, we computed assortativity, number and average size of strongly connnected components, number of edges and nodes in largest connected component, maximum node degree (in,out), number of nodes, number of edges, average node degree, and entropy of node degree sequences (In, Out). The full results for these metrics appear in SI Dataset 1 and confirm that average assortativity coefficient by sentiment is higher in the vaccine conversation (0.68) than the climate change conversation (0.44).

### Communities and their interactions

Next, we moved to the level of entire communities of users. Network communities are defined as portions of the network where individuals interact with one another much more than they interact with those outside of the community (see Methods). We measured the sizes of these communities and visualized the network of interacting communities. We also used Shannon entropy to measure how diverse these communities are with respect to user sentiment.

Vaccine communities were larger than the climate change communities, on average, for both retweets and mentions (Table 5). This was also true for the subsampled network (Table S15), the 80% threshold criterion for user classification (Table S16), and the collected dataset (Table S17). In contrast, the vaccine community diversity was higher in the vaccine discussion for the GNIP dataset under our baseline analysis (Table 5) and the GNIP dataset under subsampling (Table S15), whereas the climate change community diversity was higher for the GNIP dataset with the 80% threshold criterion for user classification (Table S16), and the collected dataset (Table S17).

**Table 5.** Vaccine conversation communities are larger and contain more diversity of user sentiment. Average community heterogeneity and size for community mutual networks using the plurality model of user sentiment. Heterogeneity is measured by Shannon entropy (see Methods).

| Average Community Heterogeneity |  |  |  |  |  |  |  |  |  |  |
| --- | --- | --- | --- | --- | --- | --- | --- | --- | --- | --- |
|  | Retweets |  |  |  |  | Mentions |  |  |  |  |
|  | 5 | 10 | 25 | 50 | 100 | 5 | 10 | 25 | 50 | 100 |
| Climate | 0.021 | 0.019 | 0.022 | 0.018 | 0.013 | 0.047 | 0.039 | 0.034 | 0.027 | 0.026 |
| Vaccine | 0.101 | 0.106 | 0.105 | 0.095 | 0.068 | 0.072 | 0.066 | 0.049 | 0.041 | 0.031 |
| Average Community Size |  |  |  |  |  |  |  |  |  |  |
| Climate | 4.125 | 4.685 | 5.866 | 7.267 | 8.950 | 2.685 | 2.894 | 3.156 | 3.217 | 3.138 |
| Vaccine | 6.692 | 7.773 | 9.836 | 11.558 | 12.449 | 3.623 | 4.352 | 5.170 | 5.385 | 5.474 |

This appears to contradict our finding on user network assortativity (Table 4), where we found that participants in the vaccine discussion interacted more often with those sharing their own sentiment across all of the permutations we explored. However, community diversity as measured by Shannon entropy is a function of both community size and makeup, when community sizes are very small. Somewhat larger communities are more likely to include individuals of two or more sentiments and thus exhibit a higher Shannon entropy. This effect will be particularly strong in this case given that the average community sizes range from 3 to 12, and given the breakdown in tweet sentiment across the three categories expressed in Table 2. The Shannon entropy increases most dramatically when a small community of individuals shifts from being homogeneous to including a single person with a different sentiment. Therefore the higher Shannon entropy of the vaccine discussion may simply reflect that the communities tend to be somewhat larger and thus more likely to include an individual with the more rare pro-vaccine or anti-vaccine sentiment, in addition to the preponderance of neutral sentiment.

To get a clearer picture of how communities of differing sentiments interact, we visualized community networks consisting of individuals tweeting 5+, 10+, or 100+ times on the topic. The climate retweet and mention community networks shows a dense network of communities dominated by pro-climate sentiment and interacting with one another (Figure 2 for communities of users tweeting 10 times or more.) Despite the non-negligible proportion of anti-climate tweets (Table 2) and users (Figure 1, SI Appendix Table S2, S3), it appears that these users are having conversations as part of a community that is dominated by pro-climate sentiment. The vaccine community networks, on the other hand, have two strongly contrasting features (Figure 2). Firstly, in the retweet network, a large community of anti-vaccine sentiment persists and interacts mostly with smaller communities that are either neutral or anti-vaccine. Similarly, pro-vaccine communities tend to interact with other pro-vaccine communities or with neutral communities. Secondly, the vaccine community network reveals a large number of isolated neutral communities. Results for for communities of users tweeting 5+ or 100+ times are qualitatively similar and appear in SI Appendix, Figures S3, S4. Overall, these network visualizations confirm a picture of much greater exchange of information between individuals of differing sentiment in the climate change conversation than in the vaccine conversation.

**Figure 2.**
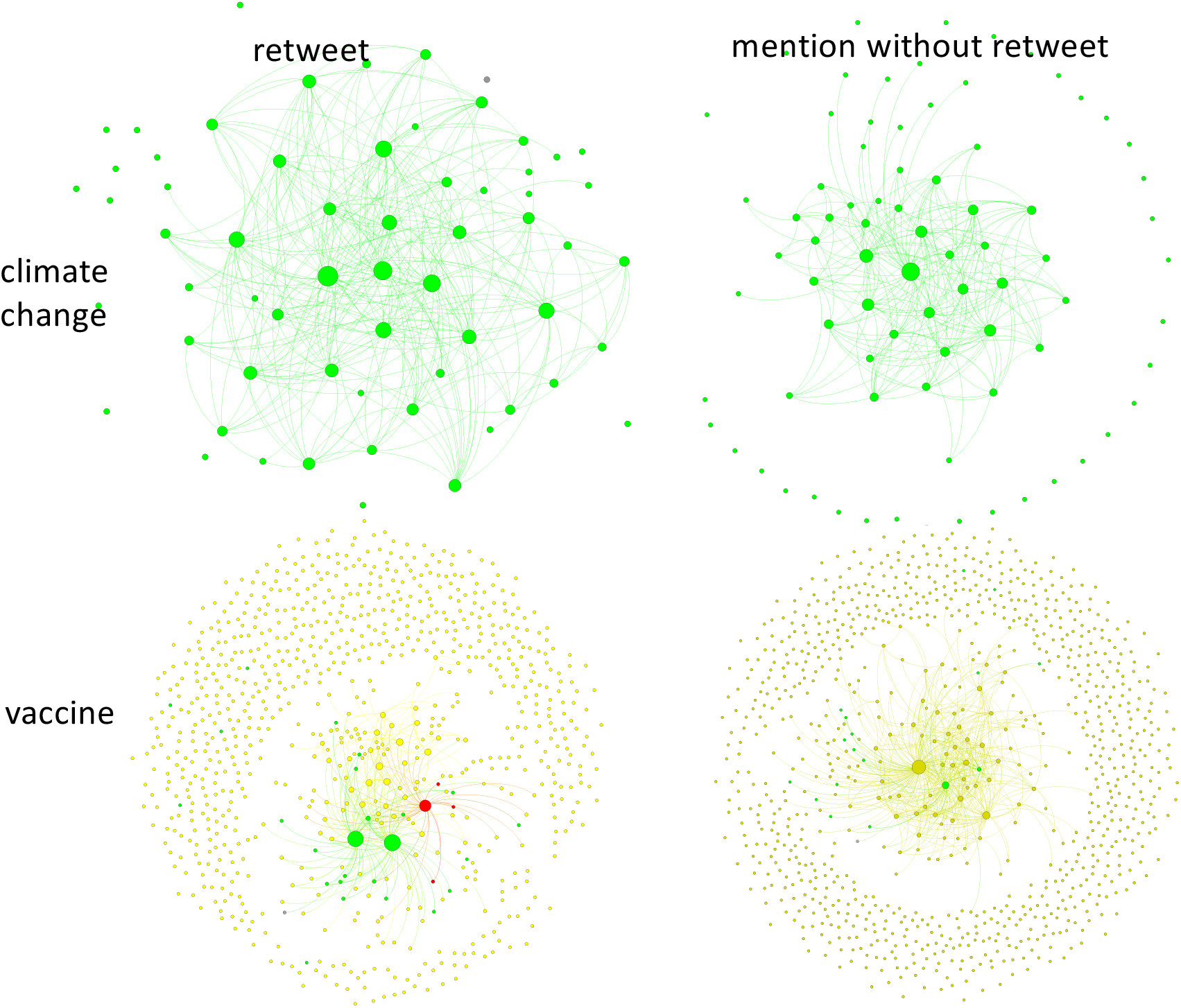
Vaccine and climate community networks exhibit distinct structure. Network of communities formed by users tweeting 10 times or more in the climate change (top) and vaccine (bottom) conversations, for retweet (left) and mention (right) networks. Each node represents a community made up of two or more users. Edges represent interactions between the users: a member of one community has tweeted or mentioned a member of another community. The size of the node represents the size of the community, and the colour of the node represents the most common sentiment of the users in each community: green represents pro-, red represents anti- and yellow represents neutral.

We note that the climate communities appear to be larger than the vaccine communities only because there are fewer of them represented in this visualization, on account of the smaller size of the climate tweet dataset. The number of communities changes with the settings on the algorithm used to determine modularity, but there are always relatively more isolated communities in the vaccine conversation than the climate conservation. For the community networks and the 260 permutations on our analysis described in the previous subsection, we also computed the number of communities, average community Size and entropy of communities (see SI Dataset 1).

## Discussion

Here we showed that individuals and communities on Twitter with differing sentiment interact significantly more in the climate change conversation than in the vaccine conversation. These two public debates share three important similarities. Firstly, both are characterized by a contrast between strong scientific consensus and divided public opinion. Secondly, both vaccine-preventable infectious disease spread and anthropogenic climate change exemplify a coupled human-environment system since there is a two-way interaction between natural processes and human processes^24^. Thirdly, both vaccine-generated herd immunity and reduced greenhouse gas emissions represent common-pool resources: they benefit everyone equally but the cost to maintain them is borne by individuals. Hence, there is a temptation to free-ride by enjoying the common resource without helping to support it^21,22,34,35^. Climate change is thought to be a particularly thorny common pool resource problem on account of the global scale of the commons involved^36^.

Qualities of real populations–such as pro-social preferences and punishment of defectors–can prevent free-riding^23,35,37^. But the temptation to free-ride suggests that we can expect to find varying sentiment in populations regarding common pool resource problems like vaccination or climate change, at least in some cases. However, despite the similarities in the systems, we found that this variation in sentiment was distributed very differently across the online social networks. We hypothesize that human systems are shaped by environmental systems in accordance to the environmental system’s spatial scale, and that this effect explains our observation. Infections spread from person to person. As a result, they typically impact local areas (except in pandemics, of course). For instance, an outbreak of hepatitis A might affect a single neighbourhood or town, and individuals may tweet about symptoms or vaccine clinics to their local networks. But climate events generates larger-scale impacts on spatial effects ranging from regions to the whole planet, and impact large groups of individuals simultaneously. This effect may explain why the vaccine community networks are characterized by a larger number of isolated communities than the climate conversation (Figure 2).

Since sentiment can also vary with geography (such as for relatively liberal coastal populations versus more conservative interior populations), the larger spatial scale of climate events may force individuals of differing sentiment to discuss climate change events more often than would occur for a natural system with localized dynamics, such as infection spread. As a result, individuals of differing sentiment may be more mixed in online social networks (Figures 1, 2). This may be a “silver lining” of a thorny global-scale common pool resource problem like climate change, but only if this increased interaction results in more widespread acceptance of anthropogenic climate change. Such a mechanism could accelerate social learning and help overcome the climate change common pool resource problem. However, this mechanism has not been explored yet in the climate change literature, either theoretically or empirically. Conversely, outbreaks of infectious diseases may improve sentiment toward vaccination, but populations that have experienced the outbreak can be expected to see the best improvement in sentiment, having experienced the negative impacts of the infectious disease firsthand. More generally, we also hypothesize that the coefficient of assortativity may indicate a community that is close to a shift social norms. Assortativity in online social media networks could therefore be monitored to track social change and perhaps provide an early warning indicator of a shift in social norms^38,39^.

Our findings include several limitations, such as classification errors in the machine learning algorithms, inability of a three-way classification system to capture nuances in sentiment, and the fact that online sentiment does not necessarily represent sentiment in the general population, or behaviour in the subpopulation that wrote the tweets. These limitations suggest opportunities for future work. For example, data on human mobility can be combined with social media data to improve our understanding of the relationship between social network structure and real-world population behaviour. Future research that expands on these aspects through comparative analysis of public conversations on vaccines and climate change may help us learn more about these highly important coupled human-environment systems.

## Methods

### Data Description

The GNIP vaccine dataset was purchased from GNIP (now, Twitter) and contains all vaccine-related tweets from April 1st, 2007 until October 15th, 2016 satisfying the search terms: Measles OR MMR OR pertussis OR DTaP OR TDap OR “chicken pox” OR (contains:vaccin) OR varicella OR “whooping cough” OR influenza OR (contains:polio) OR rotavirus OR pneumococcal OR “pneu-c-13” OR hepatitis OR meningococcal OR HPV OR flu OR (contains:immuniz) OR (contains:immunis) OR cholera OR ebola OR papillomavirus OR diphtheria OR H1N1 OR H5N1 OR H7N9 OR H3N2 OR HIV OR malaria OR mumps OR chickenpox OR rubella OR RSV OR typhoid OR (autism (jab OR (contains:shot) OR needle OR (contains:vacc) OR (contains:immuni))). The GNIP climate data set was also purchased from GNIP and contains a sample of climate change tweets from April 4th, 2015 until October 15th, 2016 satisfying the search terms: “climate change” or “global warming” (Table 1).

The collected vaccine and climate data sets (Table 1) were obtained through the Twitter Application Programming Interface (API) using a combination of the passive listener and active searches, satisfying the same search terms as the GNIP datasets. However, unlike the two GNIP datasets, the time window for both collected sets was the same, from October 16th, 2016 to August 1st, 2018. The collected dataset consists of only 2.7 and 4.4 million tweets on climate change and vaccines. The smaller size of the collected sets reflects the fact that passive and active listeners only capture a fraction of total tweets.

As reported in the Results section, we also conducted an alternative analysis that used a smaller number of more general search terms for the vaccine dataset in order to control for the generality and number of search terms used between the vaccine and climate change datasets. For this subsampling analysis we analyzed only the subset of the GNIP vaccine dataset satisfying the search terms: vaccine OR flu OR influenza.

### Tweet Sentiment Classification and Machine Learning

We created a vaccine training set starting from 75,000 randomly sampled vaccine tweets from the GNIP dataset and labelling each tweet as showing one of the following sentiments: positive, negative or neutral. Each tweet was annotated by 3 taggers on the MechTurk platform. Taggers were instructed to apply: a positive label if content of the tweet indicates that the user thinks vaccines are safe and effective; a negative label if content of the tweet indicates the user thinks vaccines are harmful and/or ineffective, or that diseases are not dangerous and do not necessitate vaccination; and a neutral label otherwise, such as when the sentiment is not clear or if the tweet sounds like a sentiment-neutral news headline and there are no additional hashtags, references, user handles, or commentaries that convey sentiment or opinion about the news article. We only included tweets in the training set where at least 2 out of 3 taggers agreed on the sentiment, resulting in 73268 tweets (Dataset 2). We used the same approach to develop a climate training set of 74895 tweets (Dataset 3). In this case, taggers were instructed to apply: a positive label if content of the tweet indicates that the user thinks climate change is real and caused by humans; a negative label if content of the tweet indicates the user thinks that climate change is not real, or that humans do not cause climate change, or that climate change is not a problem; and a neutral label for other cases, such as if the sentiment is not clear.

We also developed a third climate training set starting from 75,000 randomly sampled climate tweets from a mixture of the GNIP and collected datasets. The instructions for positive, negative, and negative sentiments were the same as above, but taggers were additionally instructed to apply a news label if the tweet sounds like a sentiment-neutral news headline and there are no additional hashtags, references, user handles, or commentaries that convey sentiment or opinion about the news article. Taggers applied a neutral label otherwise such as if the sentiment is not clear. However, for our machine learning training and all subsequent analyses, the news tweets were absorbed into the neutral class to enable comparison with the vaccine sentiment classification. Three human taggers were trained and each person tagged each tweets in the dataset. We only considered tweets where at least 2 of the taggers agreed on the sentiment, resulting in 72662 tweets. An additional dataset of 3314 negative tweets were added to supplement the initial dataset, for a total of 75976 tweets (Dataset 4).

Two machine learning algorithms were used to determine the sentiment of each tweet: a support vector machine (SVM) model and an Ensemble Averaging model. The Results section reports results using the ensemble approach unless otherwise specified. Results with both the Ensemble and SVM algorithms appear in the Supplementary Appendix.

For the ensemble approach, four models were ensembled to predict the sentiment of each tweet: a Recurrent Neural Network with Gated Recurrent units (RNN-GRU)^40^, a Recurrent Neural Network with Long Short Term Memory units (RNN-LSTM)^41^, a logistic regression classifier and a Gradient Boosted Decision Tree (GBDT)^42^. For example the Tweet ‘Climate Change is happening. We have to stop it now!’ might be scored as (0.2 negative, 0.3 neutral, 0.4 positive, and 0.1 news) by the RNN-GRU model and (0.1 negative, 0.1 neutral, 0.8 positive, and 0.0 news) by the RNN-LSTM model. The ensemble average approach determines the average score in each category, and chooses the highest scoring category as the sentiment classification. For the vaccine analysis, an equal number of tweets were randomly sampled from each sentiment category in Dataset 2 in order to create balanced training and testing sets of size of 10659. For the ensemble approach to analyzing the climate data, we used Dataset 4. We experimented with balancing the training and testing sets for this analysis but found that down-sampling the majority class (positive sentiment) ended up with a small lift in the F1 of negative sentiment prediction, but a much worse F1 in positive sentiments. Therefore, we retained use of the full, unbalanced data (Dataset 3) for the ensemble analysis of the climate change conversation (however, we note that we used a balanced training set for the SVM analysis, see next paragraph). Both training datasets were split into 80% training, 10% validation, and 10% testing subsets. The models were trained on the training set. The validation set was used for early stopping for the RNN models and pruning for the GBDT (see next paragraph). The final accuracy was determined on the testing set. For both of the RNN models, the tweets were vectorized using the glove.twitter.27B.200d pre-trained Global Vectors^43^ for Word Representation(GloVe) embeddings. The vectorized text was passed sequentially into the RNN models. The GBDT Used was lightgbm^44^, which is Microsoft’s implementation of GBDT. The RNN models were created using Tensorflow^45^. The Sklearn^46^ was used to train the logistic regression classifier. The final validation accuracy for the climate model was 78.5% on validation, with F1-scores for “Anti”, “Neutral”, “Pro”, and “News” of 0.82, 0.62, 0.83, and 0.83. The final validation accuracy for the vaccine model was 85.1% percent on validation, with F1-scores for “Anti”, “Neutral”, and “Pro” of 0.76, 0.90, and 0.77. We used the ensemble model for reporting most of our results because of the higher F1 scores compared to the SVM model.

For the SVM approach, the sentiment of each tweet was determined using support vector machine (SVM) models (sklearn version 0.19.2^46^). We again used a balanced training set generated by subsampling from Dataset 2 for the vaccine analysis. For the climate change analysis, we used a balanced training set generated by subsampling from Dataset 3. For the vaccine model the precision scores were 0.8, 0.9, and 0.79 for the negative, neutral, and positive sentiment labels respectively. The recall scores were 0.83, 0.82, and 0.82 and the F1 scores were 0.82, 0.86, and 0.81. For the climate change model the precision scores were 0.72, 0.75, and 0.58. The recall scores were 0.74, 0.51, and 0.76 and the F1 scores were 0.73, 0.61, and 0.66. Because of the variation in approaches for tagging the tweets and generating training sets for the vaccine and climate change conversations, we also conducted a separate check on the accuracy of the machine learning output by comparing the resulting classifications of user sentiment against the classifications of two human experts (see next subsection).

### User Sentiment Classification

Our objective was to classify the sentiment of users, not tweets *per se*. Hence, we also classified users into pro-/anti-/neutral on vaccines and on climate change. However, most users posted tweets belonging to more than one of the sentiment categories. The most common reasons for this were: missing contextual information (such as when a user simply retweets a headline) and errors in the automated classification by the machine learning models. To counter these sources of error, each user was labeled with a sentiment based on the most commonly tagged sentiment among their tweets (i.e., the plurality of the sentiment expressed in their tweets). Users who had an equal number of tweets in two or more of their most populous categories were labeled as undecided, although this was very rare.

To test the validity of this approach and also the validity of the machine learning classification of tweets upon which it is based, we sampled 100 randomly selected users and up to 100 randomly selected tweets from each user. The user’s sentiment based on the plurality of the automated sentiment classification of their tweets was compared with the sentiment as evaluated by a human expert using the criteria in the previous subsection to classify users into positive/negative/neutral sentiment. (Hence, a pro-vaccine user thinks that vaccines are safe and effective, for instance.) The human expert conducted their classification before seeing the results of the machine learning classification. This analysis was performed for both the GNIP climate and GNIP vaccine data sets and the Ensemble algorithm. We found that the user sentiment classification of the human expert was identical to the machine learning classification in 87% of cases for the climate dataset (meaning 13 disagreements between the machine and human expert tagged sentiment), and 84% of cases for the vaccine dataset (meaning 16 disagreements).

As reported in the main text, to assess the impact of our plurality approach to assigning user sentiment based on their tweets, we also conducted alternative analyses that classified user sentiment by requiring a threshold that 80% of a users tweets express the same sentiment in order for that user to be assigned that sentiment. For many of our analyses, we note that users were also classified into high frequency (>100 tweets), moderate frequency (>10 and < 100 tweets) and low frequency users (>5 and < 10 tweets).

### User and community networks

To study user interactions, we constructed user networks based on retweets and mentions. A retweet is when a twitter user broadcasts a tweet from another user to all of their followers. A mention is when a user includes the @ symbol followed by the user name of another user, effectively bringing the tweet to the attention of the other user. If a tweet begins with ‘RT @’ as a reply. If it appears anywhere else in the tweet then it is a mention (the user tagged with @ in the tweet will see the tweet in their mentions folder).

We considered three network classes: directed, undirected, and mutual. We built directed retweet networks from the datasets. For these networks, if A retweeted B then an edge is added from A to B. If B retweeted A then an edge from B to A is added. Each tweet corresponds to at most a single edge. The weight of the edge from A to B is the number of times A retweeted B. We built the directed mention networks in the same way, except that if A mentions (instead of retweets) B, then an edge is added from A to B. We also built undirected networks. In the undirected retweet networks if A retweeted B or B retweeted A, a single edge is added between A and B. Undirected mention networks are similarly constructed except edges are mentions instead of retweets. We built a mutual retweet networks as follows: if A retweeted B and B retweeted A a single edge is added between A and B. For the mutual mention networks if A mentioned B and B mentioned A then a single edge is added between A and B. Both types of mutual networks are also types of undirected networks, by definition. Two types of mention networks were considered: mentions with the individual being retweeted considered as a mention and mentions from non-retweet tweets only. We found that directed and mutual networks captured the full range of possible behaviours and thus we report results for directed and mutual networks only.

Network statistics were computed in Python 3.6.1 using the NetworkX package^47^ version 1.10. The networks were analyzed according to connectivity, average shortest path length, node degree, degree sequence entropy, and assortativity among other measures. We looked at the assortativity and general makeup of the networks constructed from interacting individuals. Assortativity is defined as 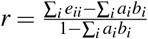. *e*_*ij*_ is the fraction of edges in a network that connect a vertex of type *i* to a vertex of type *j*. *a*_*i*_ and *b*_*i*_ represent the fraction of each type of end of an edge that is connected to a vertex of type *i*. If *r* = 1 then there is perfect assortative mixing; users are only interacting with other users who share the same opinion. If *r* = 0 then then there is no assortative mixing; users are interacting with individuals of the same or different opinion at the levels expected by the distribution of different types of individuals. If 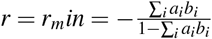, then users are interacting with those who hold differing opinions to the greatest extent permissible. Network graphs were generated in GEPHI (0.9.2-linux).

Community membership was determined using the Louvain algorithm^48^. The algorithm uses a greedy heuristic approach to optimize the modularity of a network. Modularity measures the fraction of edges within communities against the expected number of edges if the edges were distributed randomly. By optimizing modularity we identify communities with more edges within the communities than between communities. Community membership was computed using the package Python-Louvain^49^. Community networks were defined and analyzed in the same way as user networks, with results reported for mutual retweet and mention without retweet networks.

## Data Availability

The collected dataset of tweets used in this study are available from the corresponding author upon reasonable request. The GNIP dataset of tweets are available for purchase from Twitter but restrictions apply to the availability of these data, which were used under license for the current study, and so are not publicly available.

## Acknowledgements

This research was supported by a Canada Foundation for Innovation grant to C.T.B, and by a Natural Sciences and Engineering Research Council grants to C.T.B. and M.A.

## Author contributions statement

C.T.B. and M.A. conceived the study. J. Schonfeld, E.Q. J. Sinn and J.C. analyzed the data. J. Schonfeld and E.Q. generated the results. J. Schonfeld, E.Q., M.A. and C.T.B. wrote and revised the manuscript.

