## Supplementary Appendix for "Debates about vaccines and climate change on social media networks: a study in contrasts"

| Conversation | Model | Positive | Negative | Neutral | Undecided |
|---|---|---|---|---|---|
| High frequency | | | | | |
| Climate Change | support vector machine | 86.23% | 12.58% | 0.95% | 0.24% |
| Climate Change | ensemble | 97.47% | 2.08% | 0.26% | 0.18% |
| Vaccination | support vector machine | 29.32% | 10.85% | 59.61% | 0.21% |
| Vaccination | ensemble | 14.69% | 6.72% | 78.29% | 0.29% |
| Moderate frequency | | | | | |
| Climate Change | support vector machine | 86.33% | 9.48% | 2.23% | 1.96% |
| Climate Change | ensemble | 97.07% | 2.15% | 0.29% | 0.47% |
| Vaccination | support vector machine | 26.06% | 6.91% | 65.16% | 1.87% |
| Vaccination | ensemble | 14.45% | 2.64% | 81.37% | 1.54% |

Table S1: The sentiment of high frequency ($> 100$ tweets) and moderate frequency ($> 10$ tweets) users in both conversations for the GNIP datasets, according to two different machine learning algorithms.

| Conversation | Positive | Negative | Neutral | Undecided |
|---|---|---|---|---|
| High frequency | | | | |
| Climate Change | 47.9% | 7.5% | 44.2% | 0.3% |
| Vaccination | 3.9% | 7.5% | 88.5% | 0.1% |
| Moderate frequency | | | | |
| Climate Change | 58.1% | 13.0% | 25.4% | 3.5% |
| Vaccination | 4.6% | 3.6% | 91.0% | 1.0% |

Table S2: The sentiment of high frequency ($>$ 100 tweets) and moderate frequency ($>$ 10 tweets) users in both conversations for the Collected datasets, according to the ensemble machine learning algorithm.

| Threshold | GNIP_Vaccine | GNIP_Climate | Shared |
|---|---|---|---|
| 1 | 15,846,242 | 4,143,267 | 1,578,355 (9.96%, 38.09%) |
| 5 | 1,424,636 | 430,206 | 158,941 (11.15%, 36.95%) |
| 10 | 705,739 | 224,400 | 78,058 (11.06%, 34.79%) |
| 100 | 51,655 | 17,381 | 3,980 (7.70%, 22.90%) |
| Threshold | Collected_Vaccine | Collected_Climate | Shared |
| 1 | 2,219,614 | 1,356,071 | 84,985 (3.8%, 6.2%) |
| 5 | 82,396 | 59,824 | 12,761 (15.5%, 21.3%) |
| 10 | 34,665 | 23,496 | 4,328 (12.5%, 18.4%) |
| 100 | 1,307 | 572 | 78 (6.0%, 13.6%) |

Table S3: The number of unique Twitter accounts participating in the vaccine and climate change conversations thresholded by the number of tweets each account contributed to their respective data sets. The shared column contains the number of accounts that met the minimum Tweet threshold in both datasets.

| | Vaccine | | | |
|---|---|---|---|---|
| Climate | Positive | Neutral | Negative | Divided |
| Positive | 20,951 | 49,612 | 3,242 | 2,433 |
| Neutral | 11 | 274 | 10 | 3 |
| Negative | 47 | 720 | 494 | 21 |
| Divided | 12 | 154 | 70 | 4 |

Table S4: Number of users tweeting in both conversations 10 times or more on each subject, broken down by sentiment, for the GNIP dataset. Differences are statistically significant ($\chi$-squared = 3755.7, df = 9, $p < 2.2^{-16}$).

| | Vaccine | | | |
|---|---|---|---|---|
| Climate | Positive | Neutral | Negative | Divided |
| Positive | 41,462 | 98,581 | 5,416 | 8,398 |
| Neutral | 62 | 814 | 19 | 16 |
| Negative | 174 | 1,784 | 857 | 130 |
| Divided | 97 | 738 | 161 | 52 |

Table S5: Number of users tweeting in both conversations 5 times or more on each subject, broken down by sentiment, for the GNIP dataset. Differences are statistically significant ($\chi$-squared = 5871.5, df = 9, $p < 2.2^{-16}$).

| | Retweets | | | | | Mentions | | | | |
|---|---|---|---|---|---|---|---|---|---|---|
| | 5 | 10 | 25 | 50 | 100 | 5 | 10 | 25 | 50 | 100 |
| Climate | 0.996 | 0.998 | 0.998 | 0.994 | 1.000 | 0.869 | 0.896 | 0.938 | 0.947 | 0.923 |
| Vaccine | 0.995 | 0.995 | 0.996 | 0.996 | 0.996 | 0.946 | 0.964 | 0.975 | 0.977 | 0.975 |
| P 2-tailed | 0.000 | 0.000 | 0.000 | 0.000 | NA | 0.000 | 0.000 | 0.000 | 0.000 | 0.000 |

Table S6: User sentiment assortativity coefficients for the user networks: mutual network for the GNIP data using the ensemble machine learning algorithm to assign tweet sentiment and using the 80% threshold for assigning user sentiment (see Methods).

| | Retweets | | | | | Mentions | | | | |
|---|---|---|---|---|---|---|---|---|---|---|
| | 5 | 10 | 25 | 50 | 100 | 5 | 10 | 25 | 50 | 100 |
| Climate | 0.764 | 0.777 | 0.806 | 0.792 | 0.780 | 0.365 | 0.394 | 0.342 | 0.371 | 0.381 |
| Vaccine | 0.864 | 0.877 | 0.890 | 0.905 | 0.918 | 0.747 | 0.783 | 0.824 | 0.840 | 0.859 |
| P 2-tailed | 0.000 | 0.000 | 0.000 | 0.000 | 0.000 | 0.000 | 0.000 | 0.000 | 0.000 | 0.000 |

Table S7: User sentiment assortativity coefficients for the user networks: directed network for the GNIP data using the ensemble machine learning algorithm to assign tweet sentiment and using the 80% threshold for assigning user sentiment (see Methods).

| Mutual | Retweets | | | | | Mentions | | | | |
|---|---|---|---|---|---|---|---|---|---|---|
| | 5 | 10 | 25 | 50 | 100 | 5 | 10 | 25 | 50 | 100 |
| Climate | 0.380 | 0.352 | 0.308 | 0.269 | 0.226 | 0.156 | 0.139 | 0.163 | 0.210 | 0.231 |
| Vaccine | 0.824 | 0.831 | 0.839 | 0.847 | 0.856 | 0.476 | 0.475 | 0.473 | 0.463 | 0.452 |
| P 2-tailed | 0.000 | 0.000 | 0.000 | 0.000 | 0.000 | 0.000 | 0.000 | 0.000 | 0.000 | 0.000 |

Table S8: User sentiment assortativity coefficients for the user networks: mutual network for the GNIP data (and with subsampled vaccine data) using the ensemble machine learning algorithm to assign tweet sentiment and using plurality for assigning user sentiment (see Methods).

| | Retweets | | | | | Mentions | | | | |
|---|---|---|---|---|---|---|---|---|---|---|
| | 5 | 10 | 25 | 50 | 100 | 5 | 10 | 25 | 50 | 100 |
| Climate | 0.134 | 0.132 | 0.119 | 0.106 | 0.097 | 0.056 | 0.053 | 0.056 | 0.066 | 0.074 |
| Vaccine | 0.540 | 0.576 | 0.631 | 0.669 | 0.703 | 0.390 | 0.396 | 0.406 | 0.409 | 0.417 |
| P 2-tailed | 0.000 | 0.000 | 0.000 | 0.000 | 0.000 | 0.000 | 0.000 | 0.000 | 0.000 | 0.000 |

Table S9: User sentiment assortativity coefficients for the user networks: directed network for the GNIP data (and with subsampled vaccine data) using the ensemble machine learning algorithm to assign tweet sentiment and using plurality for assigning user sentiment (see Methods).

| Mutual | Retweets | | | | | Mentions | | | | |
|---|---|---|---|---|---|---|---|---|---|---|
| | 5 | 10 | 25 | 50 | 100 | 5 | 10 | 25 | 50 | 100 |
| Climate | 0.738 | 0.745 | 0.715 | 0.751 | 0.783 | 0.318 | 0.322 | 0.381 | 0.446 | 0.690 |
| Vaccine | 0.748 | 0.770 | 0.805 | 0.828 | 0.886 | 0.255 | 0.235 | 0.319 | 0.311 | 0.371 |
| P 2-tailed | 0.122 | 0.012 | 0.000 | 0.002 | 0.001 | 0.003 | 0.003 | 0.101 | 0.042 | 0.002 |

Table S10: User sentiment assortativity coefficients for the user networks: mutual network for the collected data using the ensemble machine learning algorithm to assign tweet sentiment and using plurality for assigning user sentiment (see Methods).

| Directed | Retweets | | | | | Mentions | | | | |
|---|---|---|---|---|---|---|---|---|---|---|
| | 5 | 10 | 25 | 50 | 100 | 5 | 10 | 25 | 50 | 100 |
| Climate | 0.335 | 0.340 | 0.308 | 0.319 | 0.286 | 0.203 | 0.224 | 0.218 | 0.262 | 0.286 |
| Vaccine | 0.419 | 0.502 | 0.591 | 0.652 | 0.708 | 0.280 | 0.295 | 0.374 | 0.408 | 0.447 |
| P 2-tailed | 0.000 | 0.000 | 0.000 | 0.000 | 0.000 | 0.000 | 0.000 | 0.000 | 0.000 | 0.000 |

Table S11: User sentiment assortativity coefficients for the user networks: directed network for the collected data using the ensemble machine learning algorithm to assign tweet sentiment and using plurality for assigning user sentiment (see Methods).

| | Retweets | | | | | Mention | | | | |
|---|---|---|---|---|---|---|---|---|---|---|
| | 5 | 10 | 25 | 50 | 100 | 5 | 10 | 25 | 50 | 100 |
| Climate | 0.563 | 0.552 | 0.539 | 0.514 | 0.486 | 0.276 | 0.272 | 0.301 | 0.322 | 0.341 |
| Vaccine | 0.894 | 0.902 | 0.915 | 0.922 | 0.928 | 0.699 | 0.722 | 0.746 | 0.754 | 0.749 |
| P 2-tailed | 0.000 | 0.000 | 0.000 | 0.000 | 0.000 | 0.000 | 0.000 | 0.000 | 0.000 | 0.000 |

Table S12: User sentiment assortativity coefficients for the user networks: directed network for the GNIP data using the support vector machine algorithm to assign tweet sentiment and using plurality for assigning user sentiment (see Methods).

| | Retweets | | | | | Mention | | | | |
|---|---|---|---|---|---|---|---|---|---|---|
| | 5 | 10 | 25 | 50 | 100 | 5 | 10 | 25 | 50 | 100 |
| Climate | 0.973 | 0.957 | 0.950 | 0.923 | 0.933 | 0.440 | 0.311 | 0.232 | 0.331 | 0.371 |
| Vaccine | 0.997 | 0.997 | 0.997 | 0.997 | 0.997 | 0.967 | 0.977 | 0.980 | 0.983 | 0.985 |
| P 2-tailed | 0.000 | 0.000 | 0.000 | 0.000 | 0.000 | 0.000 | 0.000 | 0.000 | 0.000 | 0.000 |

Table S13: User sentiment assortativity coefficients for the user networks: directed network for the GNIP data using the support vector machine algorithm to assign tweet sentiment and using the 80% threshold for assigning user sentiment (see Methods).

| | Retweets | | | | | Mentions | | | | |
|---|---|---|---|---|---|---|---|---|---|---|
| | 5 | 10 | 25 | 50 | 100 | 5 | 10 | 25 | 50 | 100 |
| Climate | 0.563 | 0.552 | 0.539 | 0.514 | 0.486 | 0.276 | 0.272 | 0.301 | 0.322 | 0.341 |
| Vaccine | 0.860 | 0.878 | 0.898 | 0.910 | 0.918 | 0.606 | 0.623 | 0.642 | 0.650 | 0.632 |
| P 2-tailed | 0.000 | 0.000 | 0.000 | 0.000 | 0.000 | 0.000 | 0.000 | 0.000 | 0.000 | 0.000 |

Table S14: User sentiment assortativity coefficients for the user networks: directed network for the GNIP data (subsampled for the vaccine data) using the support vector machine algorithm to assign tweet sentiment and using plurality for assigning user sentiment (see Methods).

| Avg. Community Heterogeneity | | | | | | | | | | |
|---|---|---|---|---|---|---|---|---|---|---|
| | Retweets | | | | | Mentions | | | | |
| | 5 | 10 | 25 | 50 | 100 | 5 | 10 | 25 | 50 | 100 |
| Climate | 0.021 | 0.019 | 0.022 | 0.018 | 0.013 | 0.047 | 0.039 | 0.034 | 0.027 | 0.026 |
| Vaccine | 0.194 | 0.192 | 0.202 | 0.182 | 0.171 | 0.147 | 0.137 | 0.113 | 0.089 | 0.060 |
| Avg. Community Size | | | | | | | | | | |
| Climate | 4.125 | 4.685 | 5.866 | 7.267 | 8.950 | 2.685 | 2.894 | 3.156 | 3.217 | 3.138 |
| Vaccine | 4.955 | 5.792 | 8.177 | 11.996 | 17.774 | 2.829 | 3.451 | 4.118 | 4.770 | 5.685 |

Table S15: Average community heterogeneity (measured as Shannon entropy) and size for the GNIP dataset as mutual networks using the plurality model to assign user sentiment and with subsampling of the vaccine data (see Methods).

| Avg. Community Heterogeneity | | | | | | | | | | |
|---|---|---|---|---|---|---|---|---|---|---|
| | Retweets | | | | | Mentions | | | | |
| | 5 | 10 | 25 | 50 | 100 | 5 | 10 | 25 | 50 | 100 |
| Climate | 0.002 | 0.001 | 0.002 | 0.004 | 0.000 | 0.020 | 0.013 | 0.006 | 0.006 | 0.012 |
| Vaccine | 0.001 | 0.001 | 0.001 | 0.001 | 0.001 | 0.001 | 0.001 | 0.001 | 0.001 | 0.002 |
| Avg. Community Size | | | | | | | | | | |
| Climate | 2.573 | 2.764 | 2.981 | 2.918 | 2.605 | 1.407 | 1.438 | 1.452 | 1.360 | 1.270 |
| Vaccine | 6.691 | 7.790 | 9.348 | 10.058 | 9.260 | 3.313 | 4.026 | 4.713 | 4.794 | 4.582 |

Table S16: Average community heterogeneity (measured as Shannon entropy) and size for the GNIP dataset as mutual networks using the 80% threshold approach to assign user sentiment (see Methods).

| Avg. Community Heterogeneity | | | | | | | | | | |
|---|---|---|---|---|---|---|---|---|---|---|
| | Retweets | | | | | Mentions | | | | |
| | 5 | 10 | 25 | 50 | 100 | 5 | 10 | 25 | 50 | 100 |
| Climate | 0.061 | 0.051 | 0.049 | 0.066 | 0.043 | 0.210 | 0.159 | 0.117 | 0.121 | 0.073 |
| Vaccine | 0.034 | 0.033 | 0.027 | 0.038 | 0.031 | 0.056 | 0.044 | 0.053 | 0.058 | 0.098 |
| Avg. Community Size | | | | | | | | | | |
| Climate | 1.532 | 1.627 | 1.887 | 1.982 | 1.681 | 1.817 | 1.875 | 1.872 | 1.746 | 1.355 |
| Vaccine | 2.020 | 2.420 | 3.106 | 3.705 | 3.525 | 2.154 | 2.402 | 2.693 | 2.970 | 2.857 |

Table S17: Average community heterogeneity (measured as Shannon entropy) and size for the collected dataset as mutual networks using the plurality model to assign user sentiment.

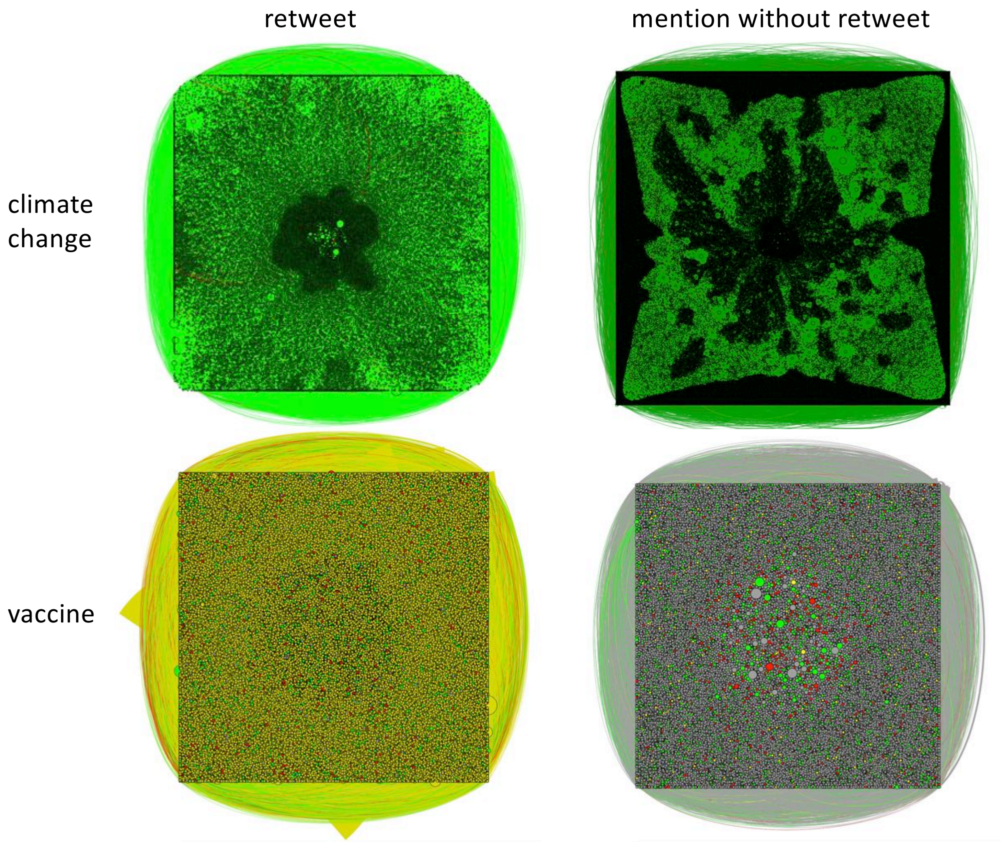


Figure S1: Network of users tweeting 5 times or more in the climate change (top) and vaccine (bottom) conversations, for retweet (left) and mention (right) networks. Each node represents an individual user. Edges represent interactions between the users: a user has tweeted or mentioned another user. The colour of the node represents the most common sentiment expressed by that user: green represents pro-, red represents anti- and yellow represents neutral.

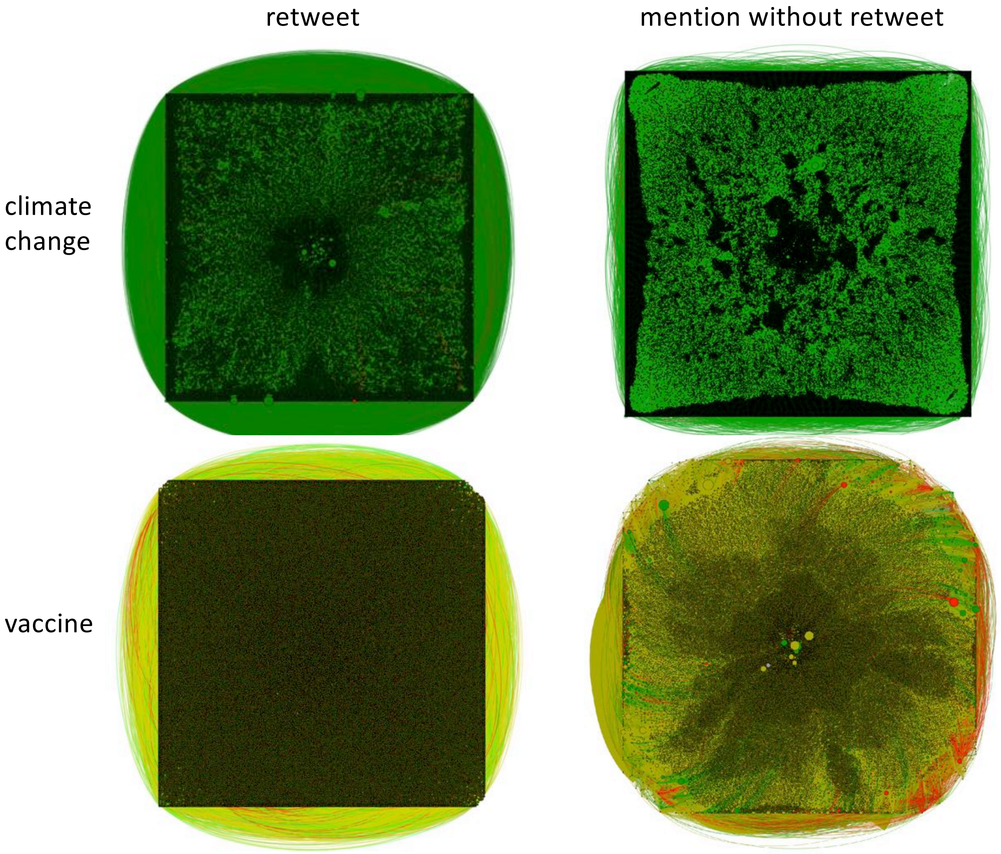


Figure S2: Network of users tweeting 10 times or more in the climate change (top) and vaccine (bottom) conversations, for retweet (left) and mention (right) networks. Each node represents an individual user. Edges represent interactions between the users: a user has tweeted or mentioned another user. The colour of the node represents the most common sentiment expressed by that user: green represents pro-, red represents anti- and yellow represents neutral.

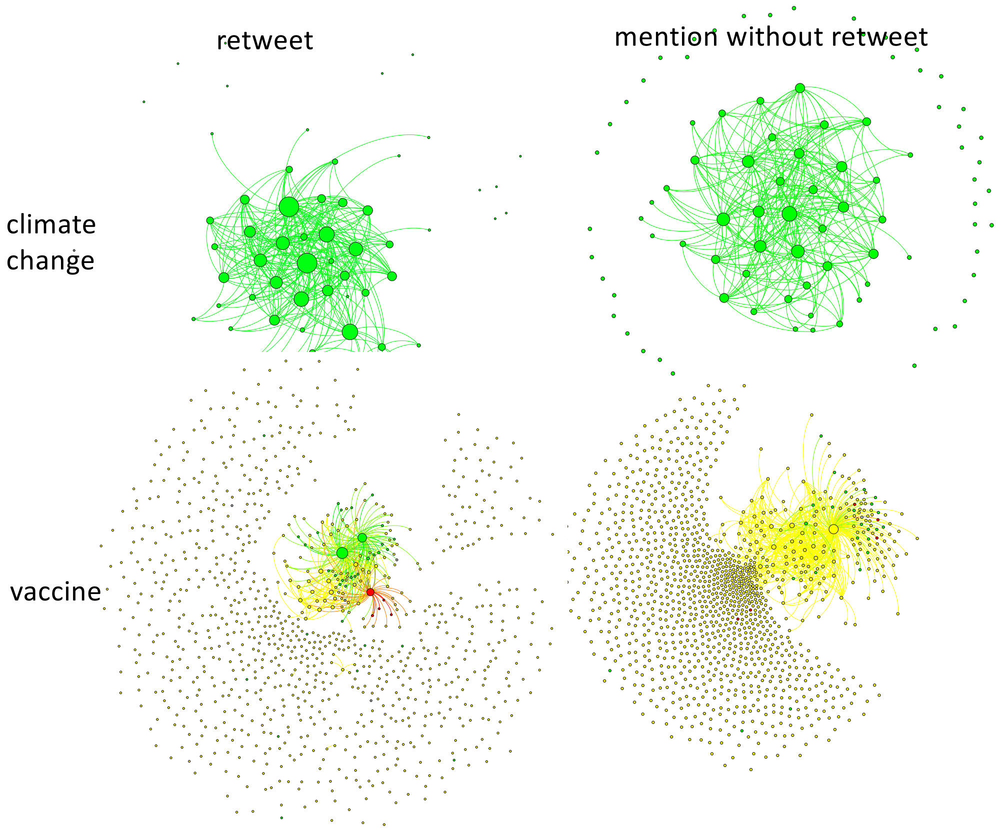


Figure S3: Network of communities formed by users tweeting 5 times or more in the climate change (top) and vaccine (bottom) conversations, for retweet (left) and mention (right) networks. Each node represents a community made up of two or more users. Edges represent interactions between the users: a member of one community has tweeted or mentioned a member of another community. The size of the node represents the size of the community, and the colour of the node represents the most common sentiment of the users in each community: green represents pro-, red represents anti- and yellow represents neutral. We note that the climate communities appear to be larger than the vaccine communities only because there are fewer of them represented in this visualization, on account of the smaller size of the climate tweet dataset.

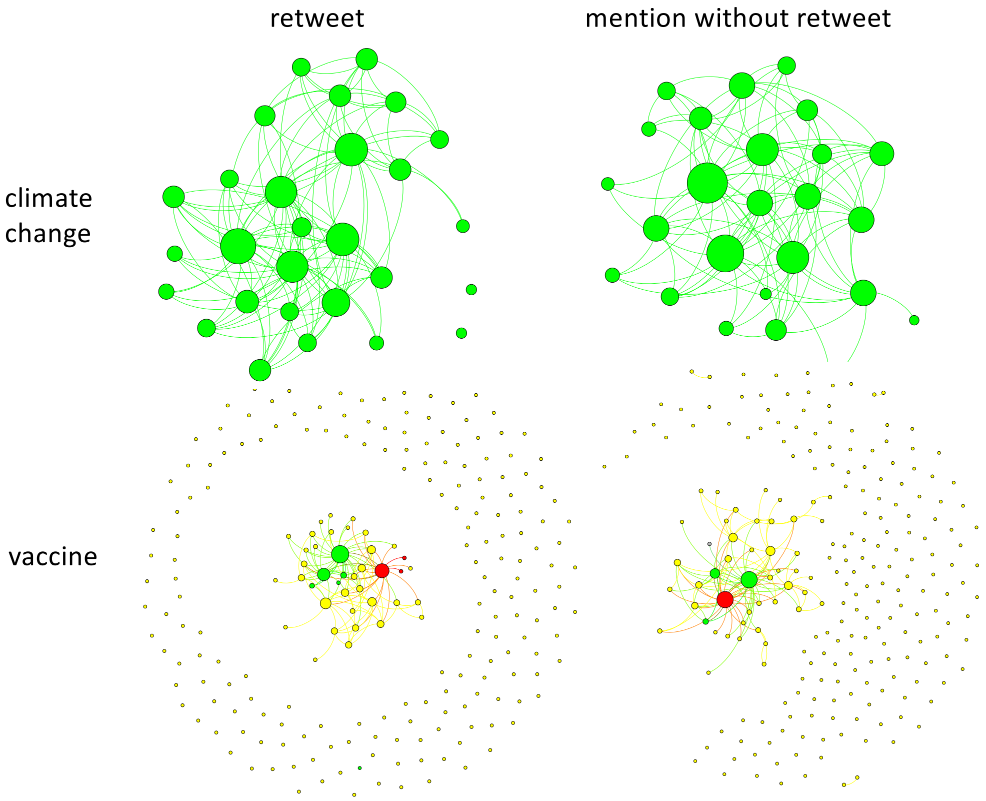


Figure S4: Network of communities formed by users tweeting 100 times or more in the climate change (top) and vaccine (bottom) conversations, for retweet (left) and mention (right) networks. Each node represents a community made up of two or more users. Edges represent interactions between the users: a member of one community has tweeted or mentioned a member of another community. The size of the node represents the size of the community, and the colour of the node represents the most common sentiment of the users in each community: green represents pro-, red represents anti- and yellow represents neutral. We note that the climate communities appear to be larger than the vaccine communities only because there are fewer of them represented in this visualization, on account of the smaller size of the climate tweet dataset.
